# HIV Envelope Trimer-Elicited Autologous Neutralizing Antibodies Bind a Region Overlapping the N332 Glycan Supersite

**DOI:** 10.1101/831008

**Authors:** Bartek Nogal, Laura E. McCoy, Marit J. van Gils, Christopher A. Cottrell, James E. Voss, Raiees Andrabi, Matthias Pauthner, Chi-Hui Liang, Terrence Messmer, Rebecca Nedellec, Mia Shin, Hannah L. Turner, Gabriel Ozorowski, Rogier W. Sanders, Dennis R. Burton, Andrew B. Ward

## Abstract

To date, immunization studies of rabbits with the BG505 SOSIP.664 HIV envelope glycoprotein trimers have revealed the 241/289 glycan hole as the dominant neutralizing antibody epitope. Here, we isolated monoclonal antibodies from a rabbit that did not exhibit glycan hole-dependent autologous serum neutralization. The antibodies did not compete with a previously isolated glycan hole-specific antibody but did compete with N332 glycan supersite broadly neutralizing antibodies. A high resolution cryoEM structure of one of the antibodies in complex with the BG505 SOSIP.664 trimer demonstrated that, while the epitope recognized overlapped with the N332 glycan supersite by contacting the GDIR motif at the base of V3, the primary contacts were located in the variable V1 loop. These data suggest that strain-specific responses to V1 may interfere with broadly neutralizing responses to the N332 glycan supersite and vaccine immunogens may require engineering to minimize these off-target responses or steer them toward a more desirable pathway.

## Introduction

Given their protective efficacy in passive transfer studies the elicitation of broadly neutralizing antibodies (bnAbs) is one of the primary objectives of current HIV research (*1–6*). Stabilized envelope glycoprotein (Env) SOSIP trimers contain all bnAb epitopes aside from the membrane proximal external region (MPER), and have provided a platform for elicitation of such bnAb responses (*7–9*). These stabilized SOSIP immunogens yield neutralization titers against immunogen-matched neutralization resistant (Tier-2) viruses in many animals (*10–16*). Previous studies (*13, 17*), and more recent imaging of polyclonal antibody responses (*18*) revealed that the primary target of neutralization on BG505 induced in rabbits is an epitope within a hole in the glycan shield. This glycan hole (GH) is mostly specific to the BG505 strain and includes a missing glycan at position N241, which is conserved in >97% of circulating Env strains (hiv.lanl.gov), and therefore there is very limited potential for broadening such responses. Bioinformatic analyses have indicated that GHs are commonly found in many HIV strains although at different positions. As illustrated by the BG505 strain, they can involve the absence of relatively highly conserved potential N-linked glycosylation sites (PNGS) (*17, 19*). Overall, these studies suggest that glycan holes are immunogenic sites that induce strain-specific neutralizing antibodies after both infection and immunization, that neutralization escape from such antibodies is relatively easy and that the corresponding responses are not on the pathway to bnAbs.

Given the limited potential of GH antibodies to develop broader reactivity, we sought to identify and characterize GH-independent neutralizing antibodies that arose in BG505 SOSIP.664-immunized rabbits (*13*). Here, we describe the high resolution cryoEM structure of a BG505 neutralizing antibody that binds to an epitope that is comprised of the variable V1 loop and that overlaps substantially with the well-known N332 glycan supersite epitope on the high mannose patch of the outer domain of gp120 (*20–22*). We conclude that these new antibodies likely do not have the potential to recognize a broad set of Envs and are potentially further problematic in their competition with bona fide bnAbs such as PGT128, PGT121, PGT135, and BG18, among others (*23–26*), making it unlikely that N332 targeting antibodies can be elicited when these V1 targeting antibodies appear.

## Results

### Trimer-elicited mAbs potently neutralize BG505 and very closely related viruses

A previous study reported the immunogenicity of BG505 and B41 SOSIP.664 immunogens in rabbits, including co-immunization with both immunogens, and serology indicated neutralizing responses outside of the GH epitope (*13*). From this cohort the post-immune plasma from 14 rabbits (Figure 1A) (*13*) were titrated against pseudoviruses derived from the MG505-BG505 mother-to-child transmission pair (*27*). Four rabbits were found to lack neutralizing activity against the previously described immunodominant glycan hole in BG505, because no gain-of-function was observed against the MG505.A2 K241S pseudovirus relative to wild type MG505.A2 in the TZM-bl neutralization assay (*13*). S241 is a critical contact residue in the center of the GH epitope that when mutated to a lysine, which naturally occurred in MG505.A2, eliminates GH-based neutralization. One animal, rabbit 5743, not only lacked GH dependent neutralization, but also had high titer neutralization activity against the wild-type MG505.A2 virus (Figure 1D). Therefore, cryopreserved primary blood mononuclear cells (PMBCs) isolated from rabbit 5743 one week after the final immunization were used to sort BG505-specific B cells. Single B cell cloning (*17*) resulted in the isolation of 12 mAbs from rabbit 5743, three of which neutralized the BG505, MG505.A2 and MG505.A2 K241S pseudoviruses, recapitulating the plasma neutralization activity of the source rabbit (Figure 1D). These three mAbs were named 43A, 43A1 and 43A2 as they are somatic variants sharing a common VDJ recombination event, although 43A2 has likely undergone a subsequent gene conversion event and thus uses a distinct VH gene.

**Fig. 1.**
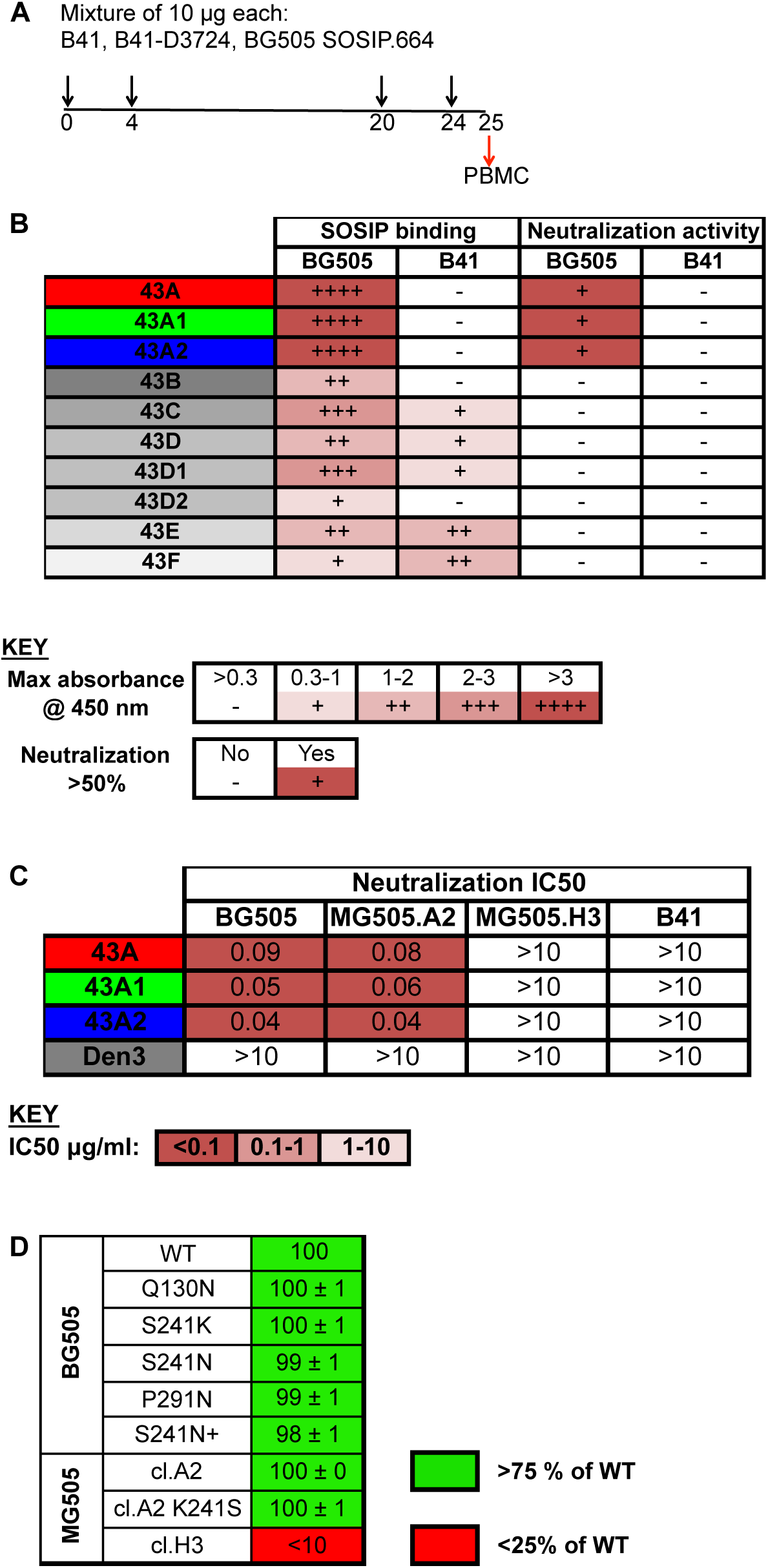
43A family antibodies neutralize BG505 and MG505 viruses. **(A)** Immunization scheme for rabbit 5743 as previously described (*13*). PBMCs were harvested 1 week following the final immunization and immunogen-specific B cells isolated by FACS as detailed in the methods. (**B**) The first two columns show a qualitative comparison (see key) of binding by 10 crude unpurified mAbs to the indicated Avi-tagged biotinylated immunogens, which were captured on streptavidin coated plates. Rabbit mAb binding was detected with alkaline-phosphatase conjugated mouse anti-rabbit IgG. The second two columns show whether these unpurified mAb preparations could neutralize the indicated pseudoviruses in the TZM-bl assay. (**C**) Purified neutralizing mAbs were titrated against the indicated pseudoviruses to generate the IC_50_ values shown. (**D**) Week 22 rabbit 5743 serum neutralization of pseudoviruses with indicated point mutations in the TZM-Bl assay (*13*).

### 43A family antibodies bind a non-glycan hole epitope

All 43A mAbs were found to bind to the gp120 subunit of the BG505 by ELISA (Figure 2A, B), but did not react with the peptide of the V3 loop of BG505 on a Fc scaffold (Figure 2C). Notably, none of the mAbs bound to the other immunogen administered during the immunization study i.e. B41 SOSIP.664 (*13*), (Figure 1A). All 43A mAbs were found to compete with one another for binding to BG505 SOSIP.664 trimer by ELISA (Figure 2D). We also did not observe competition with the previously isolated BG505 glycan hole specific mAb 11A (*17*) (Figure 2D). Moreover, the mAbs strongly competed with binding by the V3-glycan supersite bnAbs PGT121, PGT124, PGT128, 2G12, PGDM12, PGDM14 and to a lesser degree by PGT130 and PGDM21 (Figure 2D). In light of competition with these glycan-specific bnAbs, we tested the ability of the 43A antibodies to bind to a glycan array; however, no binding was detected even at high concentrations of mAb. In addition, no change in neutralization activity was observed when any of the N332 region glycan sites at positions 295, 301, 332, 386 or 392 were eliminated from the BG505 pseudovirus (Figure 2E). The potency of neutralization did however increase when the virus was grown in the presence of kifunensine, which results in relatively homogeneous Man9 glycans on the Env protein. Overall, these data suggest that the 43A mAbs primarily interact with the protein amino acids, and not the glycans themselves.

**Fig. 2.**
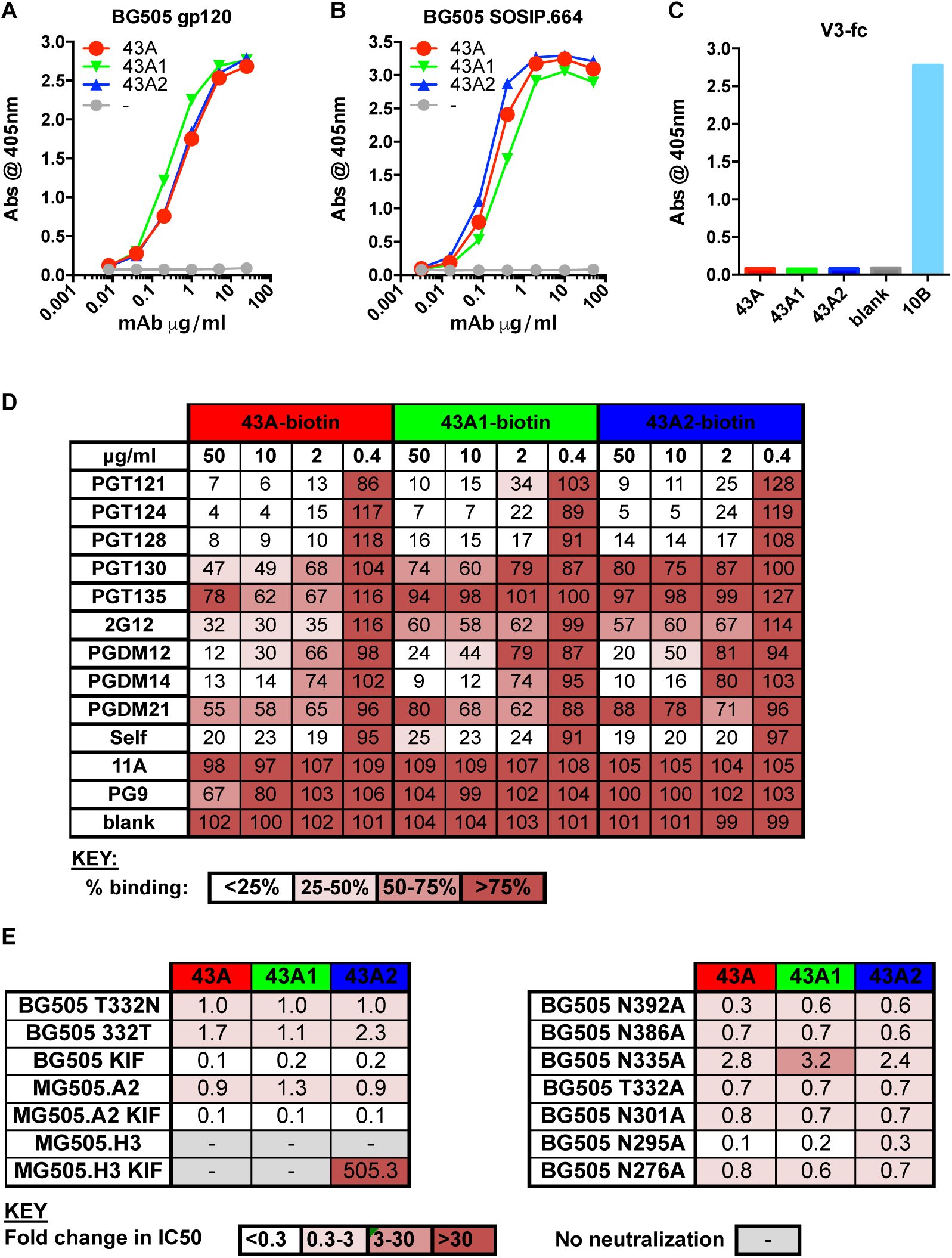
43A family antibodies bind to an epitope overlapping the bnAb N332 glycan supersite. Dose-response binding of neutralizing rabbit mAbs indicated in the legend was assayed by ELISA using streptavidin-coated plates to capture Avi-tagged biotinylated BG505 (**A**) gp120 or (**B**) SOSIP.664. (**C**) Binding by the indicated rabbit mAbs was assayed by ELISA using V3-Fc coated plates where the previously described V3 rabbit mAb 10B (*17*)was used as a positive control. (**D**) Competitor non-biotinylated mAbs listed in the right-hand column were pre-incubated with Avi-tagged BG505 SOSIP.664 protein. Binding of the biotinylated mAbs listed across the top row is expressed as percent binding, where 100% was the absorbance measured in the absence of a competitor (blank). % binding is color-coded according to the key. (**E**) The rabbit mAbs were titrated against the pseudovirus mutants indicated in the TZM-bl assay, IC_50_ values calculated using Prism and fold-change relative to the immunogen-matched strain (BG505 T332N).

### 43A mAbs neutralize MG505.A2 but not MG505.H3

Neutralization activity of the 43A mAbs was restricted to the immunogen-matched strain BG505 and the closely related MG505.A2 virus, but did not neutralize an additional virus, MG505.H3, derived from the mother. There are 13 differences in Env between MG505.A2 and MG505.H3 at positions highlighted in Figure S2. To identify which changes abrogated neutralization of MG505.H3, the 43A mAbs were tested against a panel of MG505.A2 mutant viruses where variant positions was altered to encode the MG505.H3 sequence in turn. There are 2 amino acid changes in the base of the V3 loop between MG505.A2 (G**D**IRQA**Q**) and .H3 (G**N**IRQA**H**) but altering these, either alone or in combination, had a minor (∼2.5-fold) effect on the neutralization activity of the 43A mAbs (Figure 3A). The only change that prevented neutralization by the 43A mAbs was the introduction of an extra asparagine within the V1 loop of MG505.A2 at residue 133 (CTNN) to mimic that found in MG505.H3 (Figure S2C, Figure 3A). The introduction of an alanine at the same position (CTNA) also blocked neutralization by all 3 mAbs (Figure 3A), indicating that the loop insertion rather than amino acid identity is the primary factor in escape from neutralization activity. To confirm this specificity for the V1 loop sequence in MG505.H3, two additional virus mutants were produced where the V1 loops were swapped so that MG505.A2 encoded the full V1 of MG505.H3 and vice versa. The presence of the V1 from MG505.H3 prevented all neutralization, while the presence of the V1 from MG505.A2 allowed neutralization of MG505.H3 (Figure 3A).

**Fig. 3.**
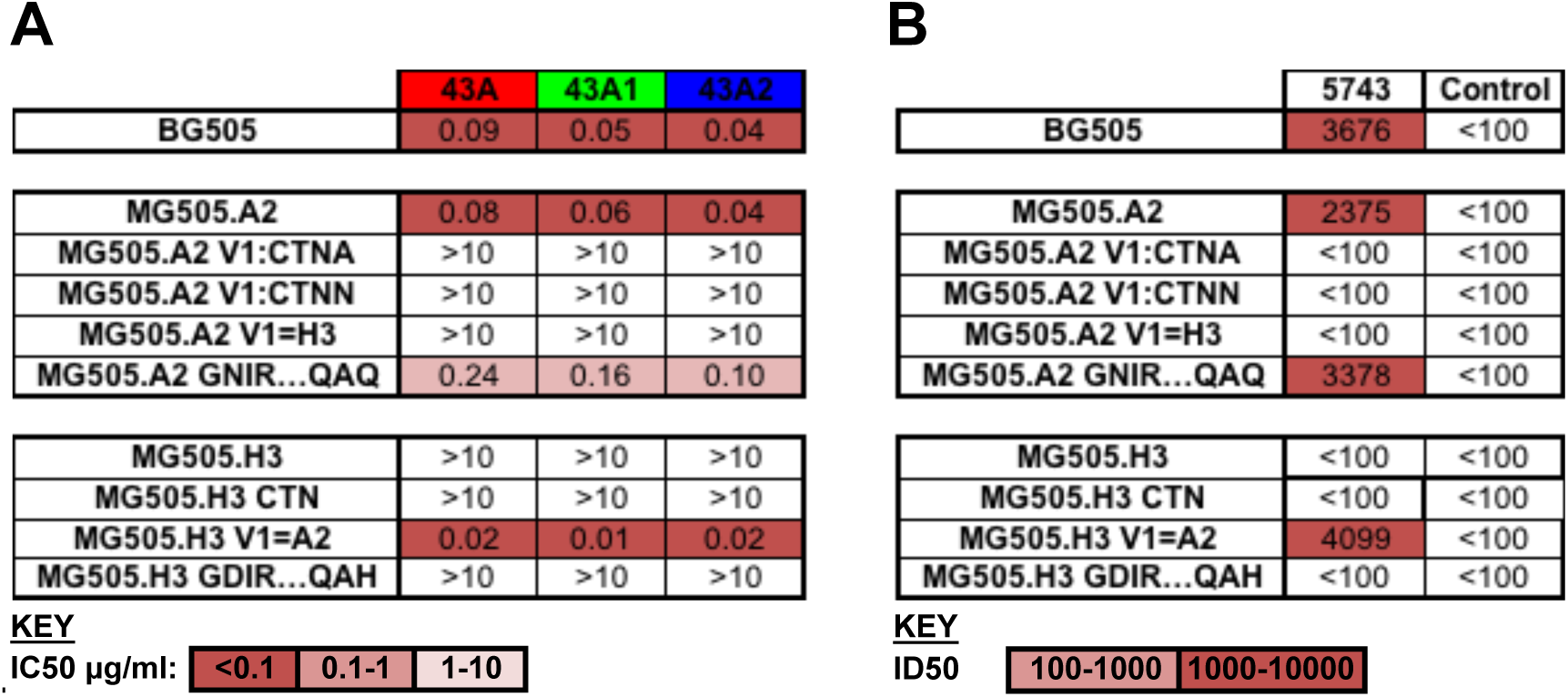
V1 loop in MG505.H3 is not compatible with 43A binding. (**A**) The rabbit mAbs were titrated against the pseudovirus mutants indicated in the TZM-bl assay, IC_50_ values calculated using Prism. (**B**) Plasma from 5743 at week 25 and from an unimmunized rabbit (Control) were titrated against the pseudovirus mutants indicated in the TZM-bl assay. 50% reciprocal dilution titers (ID_50_) were calculated in Prism. ID_50_ and IC_50_ values are color-coded according to the keys below each table.

Notably, neutralizing responses targeting the epitope bound by the 43A mAbs dominated in the source rabbit serum sample (Figure 3B) and similar serum specificities have recently been observed following non-human primates (NHP) immunization with BG505 Env (*16*). 3 of the 9 NHPs with the best BG505 neutralizing responses produced post-immune sera that could neutralize MG505.A2 but not MG505.H3(*16*). These serum samples were then tested against the V1 loop swap viruses described herein and all activity was lost when the MG505.H3 V1 loop was inserted into MG505.A2 (*16*) as seen for the 43A mAbs, although the precise epitope targeted was not revealed in this study.

### Electron microscopy studies reveal the details of 43A mAb epitopes

To confirm the epitopes identified by viral mutagenesis and binding competition assays all three 43A mAbs were digested into Fab and complexed with the BG505 SOSIP.664 immunogen, then visualized by negative stain single particle electron microscopy. The resulting 3D reconstructions revealed that all 3 neutralizing mAbs have highly overlapping epitopes near the base of V3 and similar angles of approach as previously characterized bnAbs (Figure 4A).

**Fig. 4.**
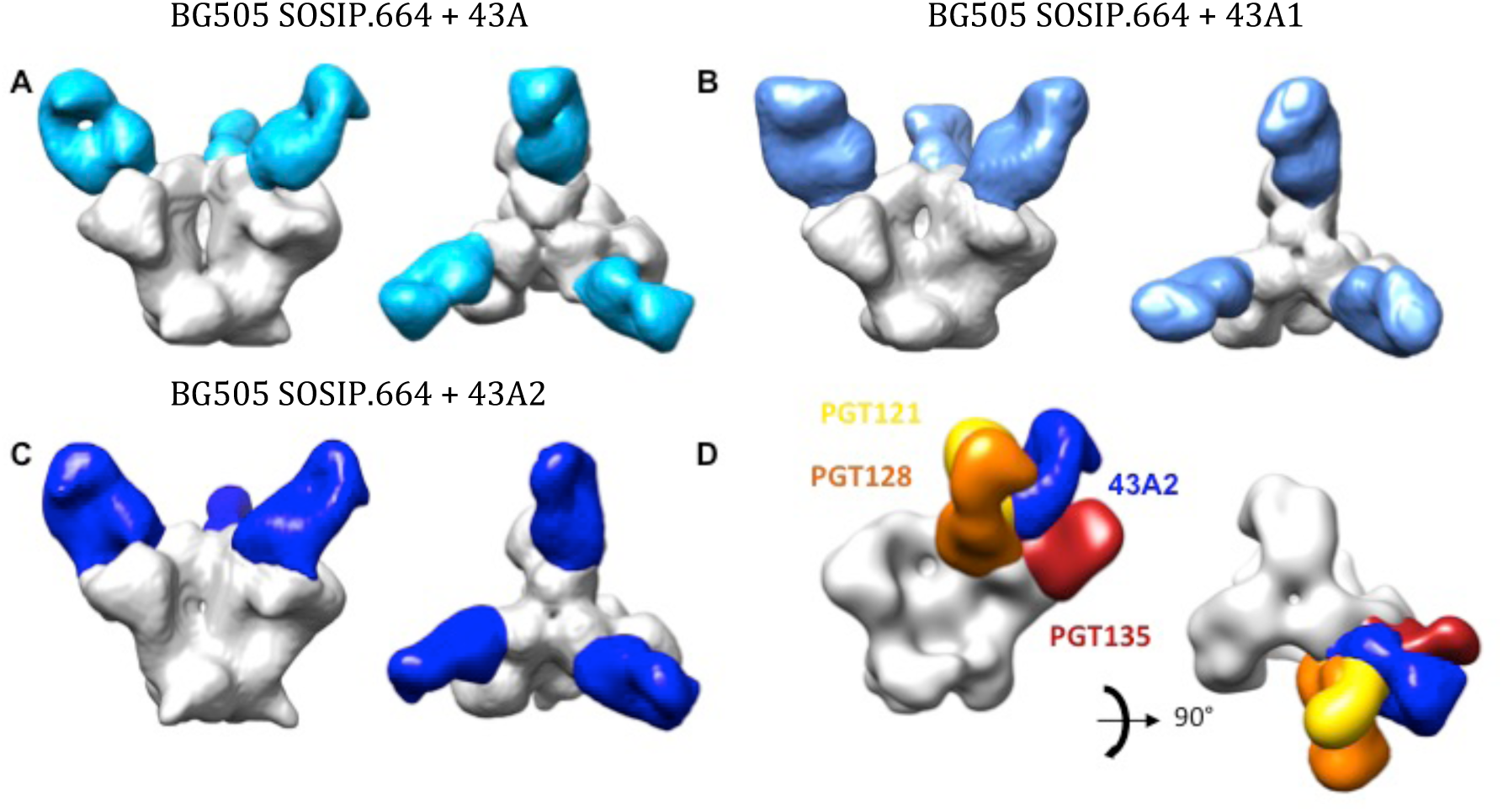
43A family antibodies all bind the gp120 at a similar angle to human bnAbs. Negative stain EM 3D reconstructions of (**A**) 43A, (**B**) 43A1, (**C**) 43A2 each in complex with BG505 SOSIP.664. The Fabs are colored in shades of blue and the Env trimer in white. (**D**) Comparison of 43A2 with N332 glycan supersite bnAbs. For clarity Fabs bound to one protomer only are shown. Side views are on the left-hand side and top views on the right-hand side of each panel.

To elucidate the molecular details of the epitopic region, the most potent antibody, mAb 43A2, was complexed with the BG505 SOSIP.664 trimer and subjected to single particle cryoEM analysis where we obtained a high-resolution reconstruction at ∼3.5 Å global resolution (Figure 5A, Table S2). This map enabled building and refinement of an atomic resolution model of the complex. The model revealed that, although the 43A2 mAb contained a 13-aa long CDRL3 that is inserted into the gp120 N332 glycan supersite region, the primary molecular contacts are with V1 of gp120. Leu94 and Asp95 of the CDRL3 make peptide backbone contacts with gp120 Ile138 and Asn136, respectively, while avoiding interaction with glycans at positions 137 and 133 (Figure 5B, S3A and S3B). In fact, there is very little interaction with any of the surrounding glycans, consistent with the neutralization data for glycan KO viruses (Figure 2D&E). Further interaction with the V1 loop is achieved via 43A2 CDRH3 Gly99 and Ser100 with the side chains of gp120 Asp140 and Asp141 (Figure S3C, D). The insertion of an amino acid in V1 resulting in loss of neutralization, such as observed in residue 133 of MG505.H3 (Figure 3A), would change the registration of the downstream V1 residues and therefore potentially disrupt interactions with Asp140 and Asp141 or result in a steric clash with the antibody.

**Fig. 5.**
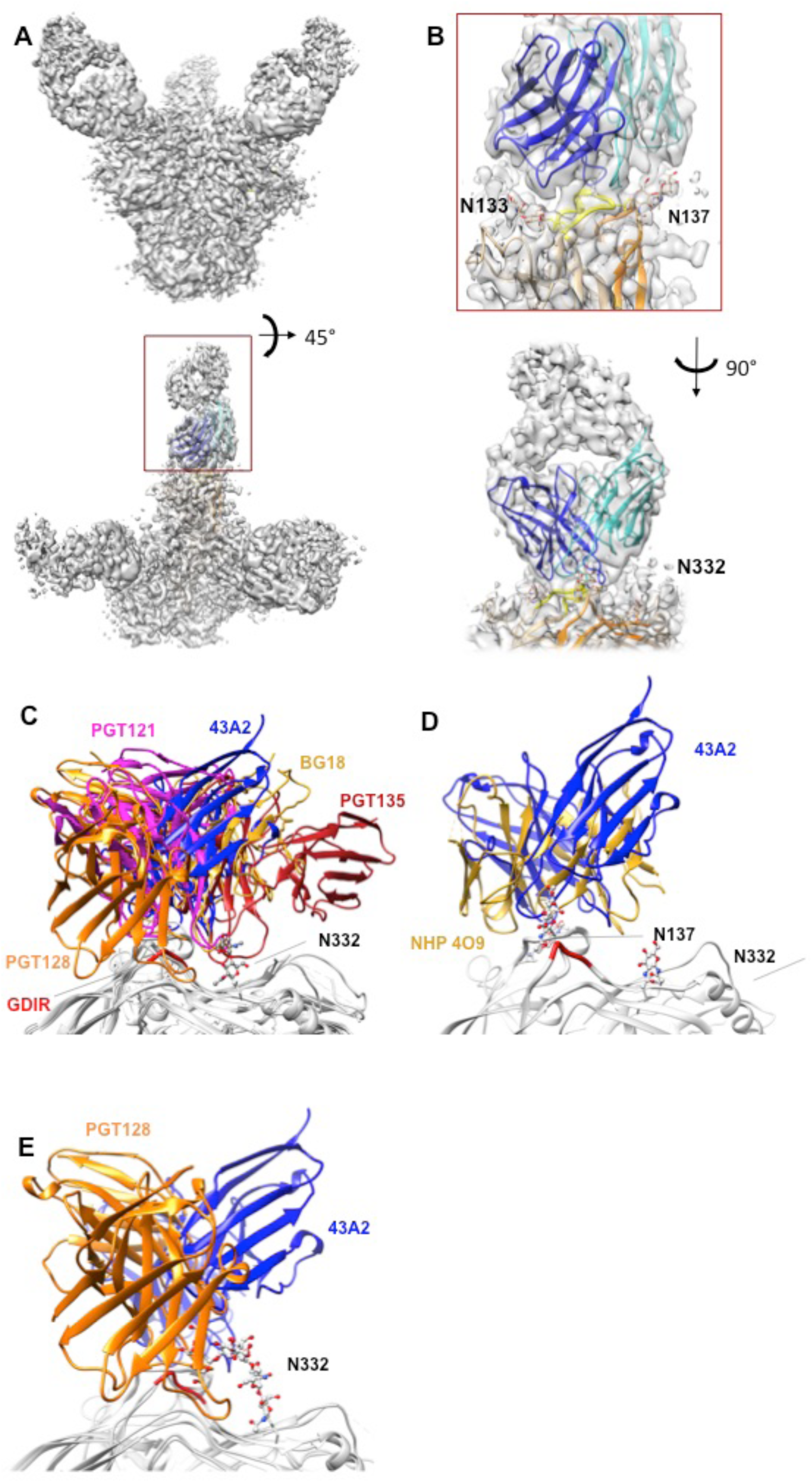
Cryo EM reconstruction of 43A2 bound to BG505 SOSIP.664. (**A**) Cryo EM map of the BG505 SOSIP.664 trimer bound three 43A2 Fabs (**B**) Chimera Modeller-generated 43A2 model showing how the antibody nestles itself within the surrounding glycans of the V1 loop (upper) as well as the N332 glycan supersite of vulnerability (lower) (**C**) Overlay with glycan supersite bnAbs PGT128 (orange, PDB: 5C7K), PGT121 (magenta, PDB: 5CEZ), BG18 (gold, PDB: 6CH7), and PGT135 (firebrick, PDB: 4JM2), showing less clashing between 43A2 and PGT135 with PGT121, PGT128, and BG18 (**D**) Overlay of 43A2 and non-human primate V1 loop binding polyclonal antibodies from animal 4O9, showing similar focus on V1 loop interaction over N332(*31*), and (**E**) Overlay of 43A2 and PGT128, showing the two share a similar footprint.

In addition to the extensive contact with V1, Leu94 of CDRL3 does contact Asp325 of the conserved gp120 GDIR coreceptor-binding motif (Figure S3E), consistent with the reduced neutralization of the GNIR mutant (Figure 3A). However, when compared to the epitopes of PGT135, PGT121, PGT128 and BG18, the bnAbs make more extensive contacts through their extended ∼20-aa CDRH3s compared to the 13-aa CDRL3 of 43A2 (Figure 5C). Notably, there is substantial overlap between 43A2 and PGT128, where both the 43A2 CDRL3 and PGT128 CDRH3 compete for the same residue 325 that is part of the GDIR motif (Figure S3E). In contrast, the weaker competition observed between PGT135 and 43A2 in the ELISA assay may be explained by the lack of engagement by the PGT135 bnAb with residues in the GDIR motif (Figure 5C).

The 43A2-bound BG505 V1 loop remains in the ground state conformation, which we define as the conformation observed in structures of Env that do not have an antibody bound to the N332 supersite or V2 apex epitopes, which influence the V1 conformation (*28–30*) The ground state conformation was also observed in the NHP V1-specific antibody revealed in a polyclonal imaging approach described elsewhere (Figure 5D and S3F) (*20, 29, 31*). In bnAb structures, the V1 loop is lifted up, providing greater access to the GDIR motif (*24, 32*). We postulate that the ground state conformation of V1 prevents additional interaction with the GDIR motif and therefore is an impediment to accessing the full extent of this important bnAb site. Further, even if the V1 loop is predisposed to an “up” conformation as inherent to N332 supersite germline targeting immunogens there are still many alternative binding poses of antibodies that likely represent off-target responses (Figure 6) (*33, 34*).

**Fig. 6.**
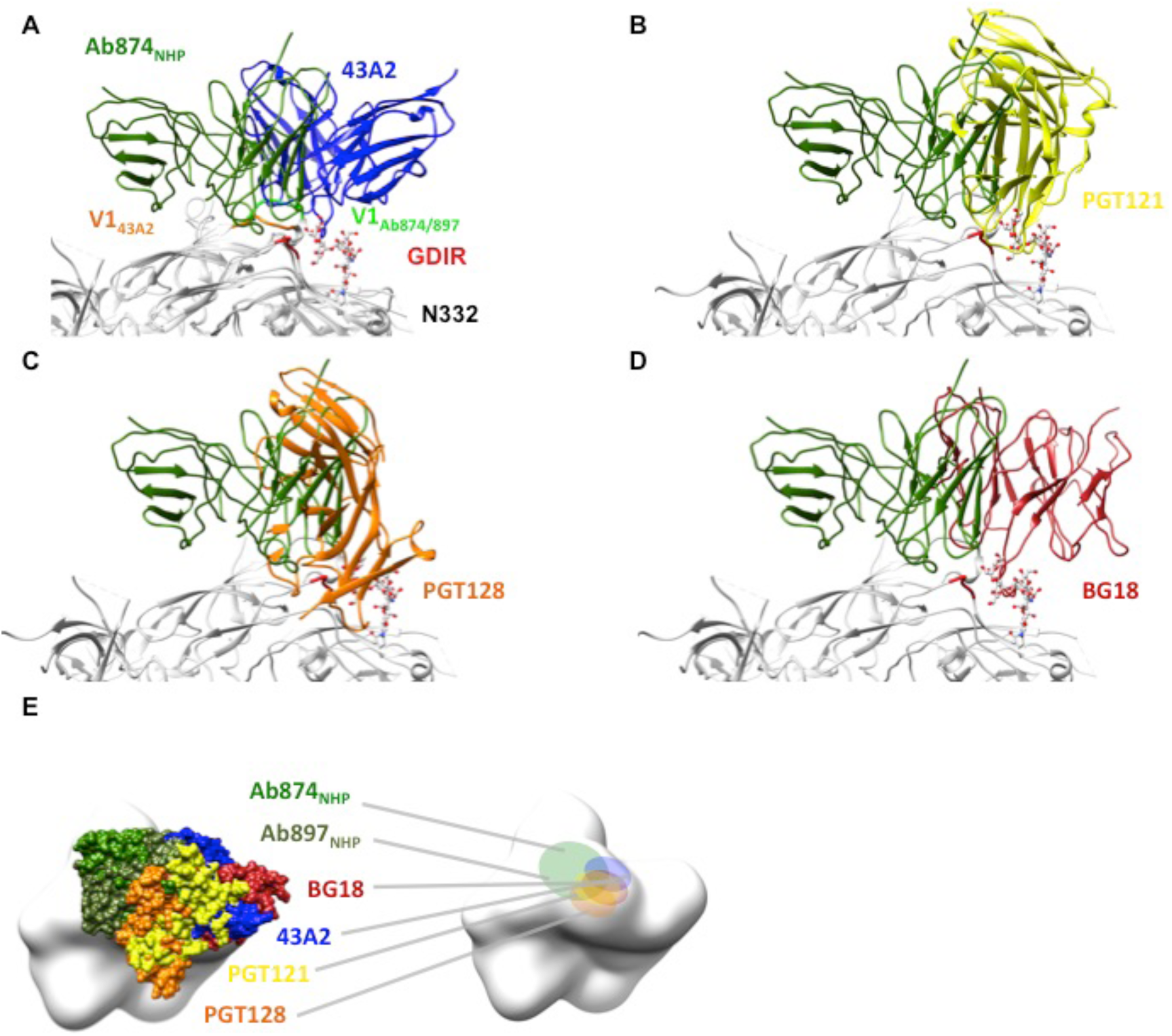
Comparison of V3 glycan-targeting bnAbs and 43A2 with non-neutralizing NHP Abs 874 and 897(*34*). (**A**) The V1 loop adopts an “up” conformation when Ab_874NHP_ and Ab_897NHP_ are bound. (**B - D**) Binding pose of Ab_874NHP_ with respect to bnAbs and (**E**) Ab_874NHP_ and Ab_897NHP_ footprint relative to bnAbs and 43A2, showing the two non-nAbs bind considerably more distant from N332 glycan supersite bnAbs.

## Discussion

Previous studies revealed that approximately 25% of rabbits immunized with BG505 SOSIP produce neutralizing responses outside of the GH epitope (*17, 35*) while, in NHPs, the number was higher (*16*). To probe details of these responses, we isolated mAbs from one such rabbit, 5743 (*13*), and subjected the Abs to antigenic and structural characterization. It is unclear why rabbit 5743 responded differently to an immunization protocol, which in three other animals produced GH-dependent responses, since all animals were co-housed and immunized at 4 months of age, limiting the potential for environmental differences. However, New Zealand White (NZW) rabbits are an outbred strain and, as such, it is plausible that genetic differences pre-disposed rabbit 5743 to react to this epitope.

Our original aim was to generate a deeper understanding of how strain-specific neutralization occurs and establish if potential for broader responses exists among these alternative epitopes. This is an important goal given the apparently limited scope to broaden the previously described GH-specific response (*17, 18, 35*). Additionally, recent studies have revealed that the glycan hole around positions 241/289 appears to be less immunodominant in non-human NHP immunization (*16*). Thus, our study describes a mAb that targets a neutralizing epitope in V1 following immunization of two different species. While the epitope footprint resembles the N332 bnAb supersite there are several key differences, and although it may be possible to broaden these V1 responses by increasing interaction with the more conserved GDIR motif, there are several hurdles that need be overcome. Thus, we conclude that the V1 is a hotspot for strain-specific antibodies that are likely to compete with the N332 bnAb supersite epitope.

Current understanding of how human bnAbs bind the glycan supersite suggests there are two ways neutralizing antibodies attack this area to achieve breadth: (1) bind directly to conserved glycans, such as N332, and (2) bind to the conserved GDIR motif by inducing a conformational change in the V1 loop that typically restricts access to the site. PGT128, PGT121 and BG18 are examples of bnAbs that employ both mechanisms (*20, 21, 26*). In contrast to bnAbs, the neutralization activity of the rabbit 43A mAbs is not adversely affected by the removal of any particular glycan, nor is there evidence of specific glycan contacts in our cryoEM map. Furthermore, virus produced with artificially homogenous Man9 glycans is more potently neutralized, which, along with our cryoEM data, suggests that the underlying peptide is better exposed when glycans are more homogeneous. Thus, while the 43A mAbs have effectively navigated the glycan shield near the high mannose patch on the gp120 outer domain they are biased toward the variable V1 loop and are not capable of providing broad neutralization.

Given our results, we conclude that elicitation of bnAbs against the N332 glycan supersite using wild-type Env sequences remains a substantial challenge. Strain-specific nAbs do not typically include bnAb features like long CDRH3 loops, primarily bind to epitopes comprised of peptide only, present lower bars to elicitation than bnAbs and can be highly competitive with bnAbs. Thus, immunogens that are designed to exquisitely stimulate appropriate precursors and then shepherd responses along narrow pathways to bnAbs may be required. Indeed, such an approach has been shown effective in heavy chain knock-in mice with heavily biased precursor repertoires (*36, 37*). However, in more realistic situations, such as human vaccination, involving greater competition between antibody responses, we hypothesize that the achievement of broad neutralization will be substantially more challenging. Of note, a recent study reported the elicitation of antibodies to the N332 supersite epitope in wild-type mice and NHPs (*34*). However, despite biasing the N332 epitope by removing a conserved glycan (N156) from the original germline-targeting (GT) immunogen (*36, 37*), the elicited antibodies were not neutralizing. Structures of these three antibodies (*34*) in complex with the cognate immunogens revealed some weak similarity to the recognition mode of bnAbs, but the antibodies lacked the hallmarks of bnAbs including a long HCDR3 and explicit recognition of conserved glycans (Figures 6 and S4; Table S1), similar to nAb 43A. Interestingly, the antibodies did bind with the V1 in the up conformation (Figure 6A, S4D), which is likely predisposed to this conformation by the N136P mutation (Fig. S4D) introduced in the original GT trimer (*36, 37*). However, due to the removal of glycans employed in the immunogen design, the antibodies were located relatively far from the core of the bnAb epitope, even farther than the 43A nAbs43A2 nAb (Figure 6B-E). The NHP antibodies also made specific contacts with several introduced mutations as well as the N156Q mutation introduced to remove the glycan (Table S1). It is therefore difficult to see how the NHP antibodies, and their corresponding binding pose, and contact with N156Q, could be further matured to recognize an intact glycan shield and cross-react with a large diversity of Envs. Thus, glycans play a key role in steering immune responses toward the correct footprints and binding poses.

Overall, although it appears possible to target features of the N332 supersite in different models (Table S1) using available immunogens, substantial hurdles remain to eliciting authentic bnAbs to this site. Although these strain-specific nAbs target an overlapping epitope the details of the molecular interactions are quite different, and more worryingly, they can compete with Abs that may have higher probability to evolve into broad neutralization, and thereby potentially suppress bnAb responses.

## Materials and Methods

### Experimental Design

Rabbit 5743 serum exhibited strong autogous neutralization of the BG505 pseudovirus, but unlike neutralizing sera of most rabbits, 5743 also neutralized the closely related MG505.A2 strain, where a lysine at position 241 abrogates GH neutralization. Thus, the objective of this study was to characterize and map the response(s) exhibited by rabbit 5743 serum. Toward that end, we isolated PBMCs and pefromed BG505-specific B cell sorting to isolate the relevant mAbs. Subsequently, site-directed mutagenesis of pseudoviruses and ELISAs were used to approximate the BG505 epitope to which the 5743 neutralizing were elicited, and whether these isolated mAbs overlap with known bnAbs. In order to deteremine the degree to which glycans were involved in the paratope-epitope interactions, we also performed neutralization assays with deglycosylated BG505 and MG505.A2 pseudoviruses. To visually confirm the epitope of the 43A class of mAbs, negative-stain EM was used, followed by high-resolution CryoEM with 43A2 in order to elucidate the details of the BG505 epitope-43A2 paratope toward informing immunogen design.

#### Isolation of rabbit B cells

Cryopreserved PBMCs from rabbit 5743 were thawed, resuspended in 10 ml of RPMI 10% FCS, and collected by centrifugation at 600 × g for 5 min. Cells were washed with PBS and resuspended in 10 ml of PBS and collected by a second centrifugation step. Cells were resuspended in 100 *μ*l of FWB (2% FCS PBS) with anti-rabbit IgM FITC (1:1000) and a streptavidin-APC tetramer of biotinylated anti-rabbit IgG. After 1 h on ice, cells were washed once with 10 ml of PBS, collected by centrifugation, and resuspended in 100 *μ*l of FWB with 1 *μ*l of a streptavidin-PE tetramer of biotinylated BG505 or B41 SOSIP.664. After a further 1 h on ice, cells were washed once with 10 ml of PBS, collected by centrifugation and resuspended in 500 *μ*l of FWB for sorting on a BD FACS Aria III. IgM-IgG+BG505+B41 lymphocytes were collected at 1 cell per well into Superscript III Reverse Transcriptase lysis buffer (Invitrogen) as previously described and immediately stored at −80°C prior to cDNA generation and single cell PCR.

#### Generation of antibodies and Fabs

Rabbit Ab variable regions (Genbank accession numbers: KX571250-1324) were cloned into an expression plasmid adapted from the pFUSE-rIgG-Fc and pFUSE2-CLIg-rK2 vectors (Invivogen). Human and rabbit Abs were transiently expressed with the FreeStyle 293 Expression System (Invitrogen). Abs were purified using affinity chromatography (Protein A Sepharose Fast Flow, GE Healthcare) and the purity and integrity were checked by SDS–PAGE. To generate Fabs, rabbit IgG was digested with 2% papain (Sigma P3125) in digestion buffer (10 mM L-cysteine, 100 mM Na acetate pH 5.6, 0.3 mM EDTA) for 6 h then quenched with 30 mM iodoacetamide. Undigested IgG and Fc fragments were removed by affinity chromatography and the Fab-containing flow through was collected. Size-exclusion chromatography was performed using Superdex 200 10/300 resin (GE Healthcare) to remove papain and digestion byproducts.

#### Neutralization assays

Pseudovirus neutralization assays using TZM-bl target cells were carried out as previously described (*38*). Briefly, single-round infectious HIV Env pseudoviruses were made as described previously (*39*). Briefly, plasmids encoding Env were cotransfected with an Env-deficient backbone plasmid (pSG3DENV) using Fugene 6 (Promega). Virus-containing supernatants were harvested 48 hr post-transfection, stored at −80°C and then titrated on TZM-bl target cells to determine the dilution needed for neutralization assays. Filter-sterilised mAbs and/or plasma were then serially-diluted in a 96-well plate and incubated with virus for 1h prior to the addition of TZM-bl target cells. After 48 hours the relative light units (RLU) for each well were measured and neutralization calculated as the decrease in RLU relative to virus only control wells. ID50/IC50 values reported as the reciprocal dilution/antibody concentration that resulted in 50% virus neutralization after fitting the curve of log concentration (plasma/mAb) versus percent neutralization in Prism. For kif-grown viruses, 25 mM kifunensine was added to 293T cells on the day of transfection. To produce mutant viruses the indicated Env encoding plasmid (BG505, MG505.A3 or MG505.H3) was altered by site-directed mutagenesis using the QuikChange site-directed mutagenesis kit (Agilent) according to the manufacturer’s instructions. Sanger sequencing was performed to verify that each plasmid encoded the desired mutation. Mutant pseudoviruses were then produced by co-transfection with pSG3DENV as described above.

#### Enzyme-linked immunosorbent assay (ELISA)

ELISA assays were performed as previously described (*40, 41*). Binding of rabbit mAbs was assayed by ELISA using streptavidin-coated plates to capture avi-tagged biotinylated antigen, namely BG505 or B41 SOSIP.664. 96-well plates were coated overnight at 4°C with streptavidin (Jackson Immunoresearch) at 2 μg/ml in PBS. Plates were washed 4 times with PBS, 0.05% (v/v) Tween, and blocked with 3% (w/v) BSA PBS for 1 h. Subsequently, 1 μg/ml of purified antigen (specifically biotinylated via a C-terminal Avi-tag) was added for 2 h. Plates were washed four times and incubated with serial dilutions of rabbit mAbs for 1 h (either crude preparations or purified as indicated in the figure legends). Plates were then washed again and binding detected with anti-rabbit Fc conjugated to alkaline phosphatase (Jackson Immunoresearch) at 1:1000 for 1 h. For gp120 protein and the V3-FC construct proteins were directly coated onto the ELISA plates and the 2 h antigen capture step omitted.

#### Competition Enzyme-linked immunosorbent assay (ELISA)

96-well ELISA plates were coated overnight at 4°C with mouse anti-Avi-tag antibody (Genscript) at 2 μg/ml in PBS. Plates were washed 4 times with PBS, 0.05% (v/v) Tween, and blocked with 3% (w/v) BSA PBS for 1 h. Concurrently, 5-fold serial dilutions of non-biotinylated rabbit or human mAbs starting at 50 μg/ml were pre-incubated with 1 μg/ml of purified Avi-tagged BG505 SOSIP.664 protein for 1 h. The mAb-SOSIP mixture was then transferred to the blocked ELISA plates and incubated for 1 h. Plates were washed four times and incubated with 0.5 μg/ml of biotinylated mAb for 1 h, then washed again and binding detected with streptavidin-alkaline phosphatase (Jackson Immunoresearch) at 1:1000 for 1 h. mAbs were biotinylated using the NHS-micro-biotinylation kit (Pierce). Competition is expressed as percentage binding where 100% was the absorbance measured when BG505 SOSIP.664 protein only was captured on the anti-Avi-tag ELISA plate.

#### Negative-Stain EM Sample Preparation

43A, 43A1, and 43A2 Fabs and BG505 SOSIP.664 (± D368R/N276A) trimers were expressed in 293F cells and purified using a previously described procedure (*42*). Briefly, the trimers were affinity purified using 2G12 antibody resin, buffer exchanged into 20 mM Tris; 0.5M NaCl, pH 8.0 (TBS), followed by removal of aggregates, monomers, and dimers via size exclusion chromatography (SEC). 43A2 IgG was purified by MaSelect Protein A, then digested into Fab using papain resin, followed by further purification with Protein A in order to remove the Fc domains. Trimer-Fab complexes were formed using a range of 6-molar excess of Fab-to-trimer

#### Negative-Stain Electron microscopy

At a concentration of ∼0.03 mg/mL, the Fab-trimer mix was applied to glow-discharged, carbon-coated 400 mesh copper grids, followed by blotting to remove excess sample. 3 μlμl of 2% (w/v) uranyl formate stain was applied and blotted off immediately, followed by application of another 3 μl of stain for 45-60s, again followed by blotting to remove excess stain. Stained grids were allowed to air-dry and stored under ambient conditions until ready for imaging. Images were collected via Leginon software (*43, 44*) using the Tecnai T12 electron microscope operated at 120kV x 52,000 magnification. The electron dose was 25 e^−^/A^2^. Particles were picked from the raw images using DoG Picker (*45*) and placed into stacks using Appion software (*46*). Initial 2-D reference-free alignment was performed using iterative multivariate statistical analysis/multi-reference alignments (MSA/MRA) to generate a relatively clean stack of particles (*47*). Next, the clean particle stacks were converted from IMAGIC to RELION-formatted MRC stacks and subjected to RELION 2-D classification (*48*), followed by RELION 3-D reconstruction.

#### CryoEM Sample Preparation

500 μg of trimer was incubated with tenfold molar excess of 43A2 Fab overnight. The complex was then purified over Superose 6 column (GE Healthcare) and concentrated to 1.5 mg/mL and mixed with Lauryl Maltose Neopentyl Glycol (LMNG, Anatrace) prior to deposition onto 2/2 Quantifoil grids (EMS) that were glow discharged for 10 s directly preceding the deposition in a Vitrobot (Thermo Scientific). Once sample was deposited the grids were blotted and plunged into liquid ethane using the Vitrobot to immobilize the particles in vitreous ice. Using Leginon image acquisition software, we collected 1,366 micrographs at a nominal magnification of 29,000x with a Gatan K2 summit detector mounted on a Titan Krios set to 300 kV set to counting mode for the data collection(*44*). The dose rate was ∼4.78 e^−^/pix/s with frame exposure of 250 ms, with a total exposure time and dose of 14 s and 60 e^−^/Å^2^, respectively. MotionCor2 was used for frame alignment and CTF models were determined using GCTF (*49*). DogPicker was used to pick 455,207 particles, which were then extracted and subsequently 2D-classified in cryoSPARC (*45, 50*). Selected 2D classes amounting to 85,841 particles were then fed into the 3D homogeneous refinement algorithm using C3 symmetry, resulting in final resolution of ∼3.52 Å.

#### Model Building and Refinement

The BG505 SOSIP.664 trimer structure from PDB 5ACO was docked into the cryoEM map using UCSF Chimera (*51*). Subsequent iterations of manual and Rosetta fragment library based centroid rebuilding and refinement were then performed (*52*). The resulting model was then refined using all atom refinement under constraints of the density map. Glycans were manually built in Coot (*53*). 43A2 model was generated using Chimera Modeller, fit into the cryo EM density and subsequently manually built in Coot, followed by a final Rosetta refinement (*53, 54*).

## Acknowledgements

We are grateful to Bill Anderson for expert microscopy assistance. The research was supported by NIH grant UM1AI100663 (ABW and DRB), UM1AI144462 (ABW and DRB) and P01 AI110657 (ABW and RWS), the Bill and Melinda Gates Foundation grants OPP1115782 (ABW) and OPP1132237 (RWS), amfAR grant 109514-61-RKVA (MJG). RWS is a recipient of a Vici grant from the Netherlands Organization for Scientific Research (NWO). C.A.C. was supported by NIH F31 Ruth L. Kirschstein Predoctoral Award Al131873 and by the Achievement Rewards College Scientists Foundation.

---

**Fig. S1. Related to Figure 1.**
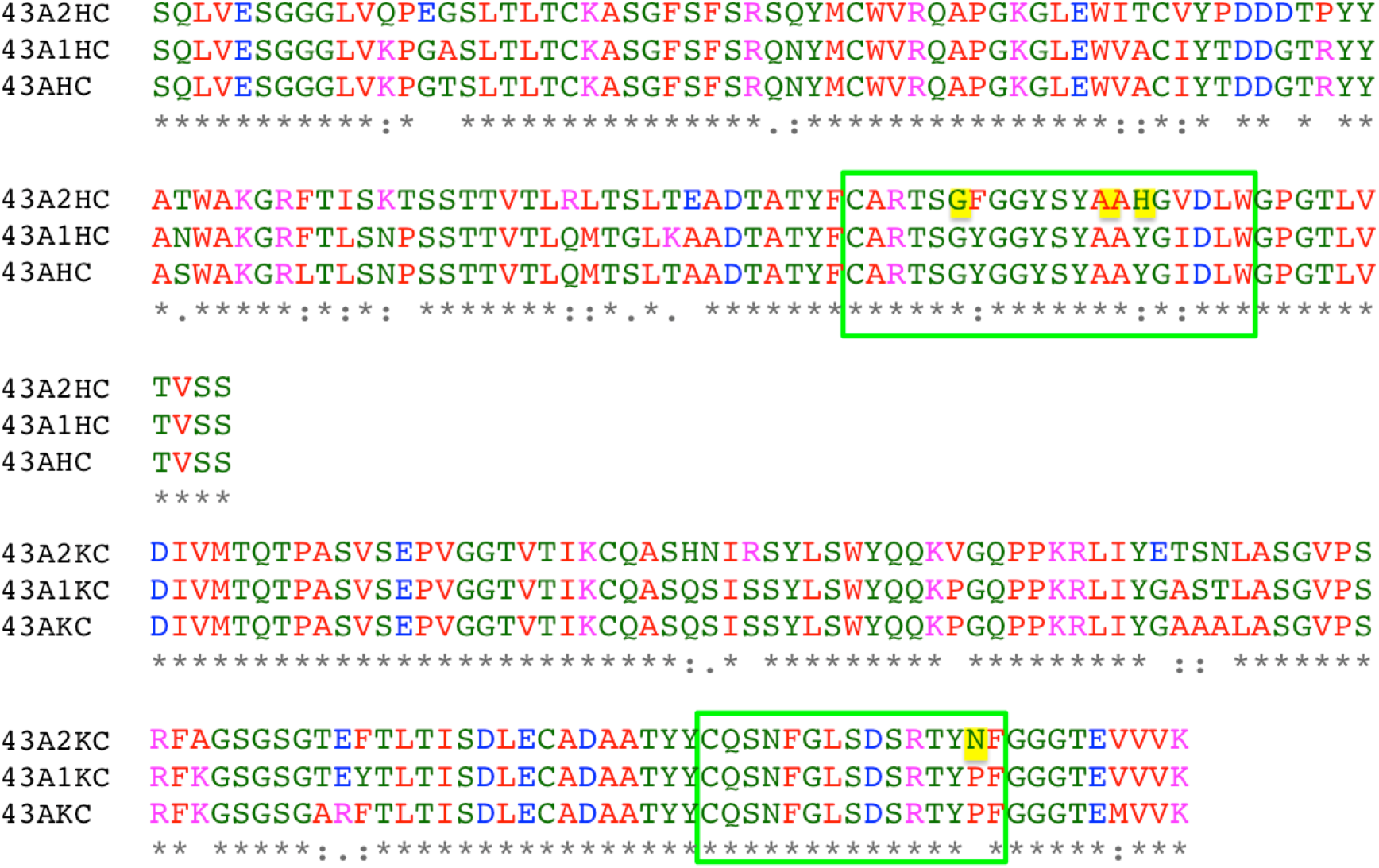
Sequence alignment of the heavy and light chains of the 43A monoclonal antibodies.

**Fig. S2.**
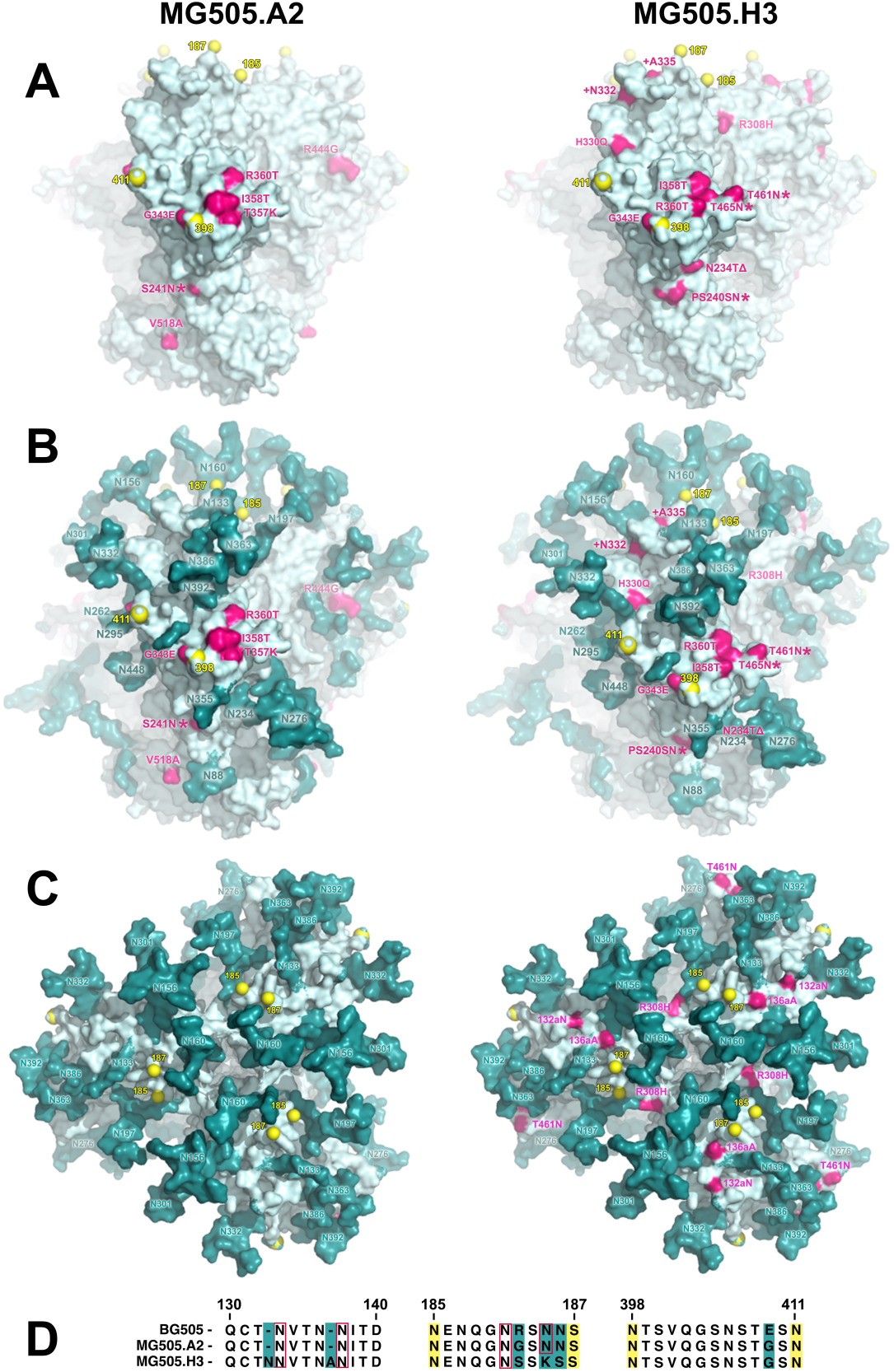
Comparison of MG505.A2 and MG505.H3 Env. (**A**) MG505.A2 (left-hand panel) and MG505.H3 (right-hand panel) amino acid sequences modelled onto the structure of crystal structure of BG505 SOSIP.664 (PDB: 53TX). Residues colored pink differ between MG505.A2 and BG505 (left-hand panel) and between MG505.H3 and BG505 (right-hand panel). Residues colored yellow mark the limit of the predicted structures (between yellow spheres are unstructured loops). (**B**) Same models as in (**A**) but with N-linked glycans are highlighted in teal and numbered. (**C**) Apex view of the structures shown in (**B**). (**D**)An alignment of sequence changes between MG505.A2 and MG505.H3.

**Fig. S3. Related to Figure 5.**
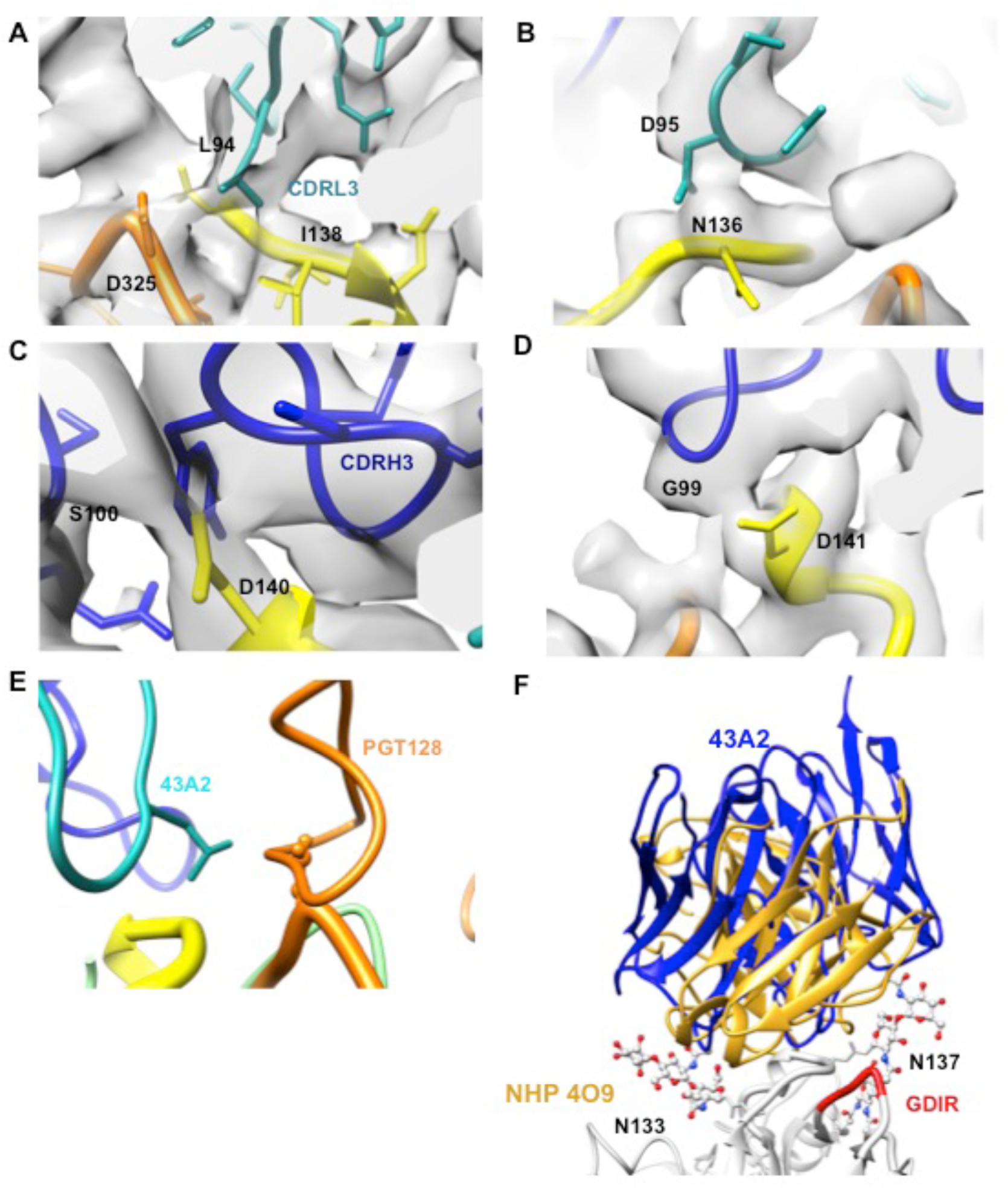
CryoEM map details of the 43A2–BG505 SOSIP.664 interactions. (**A**) and (**B**) CDRL3 interactions with the V1 and V3 loops, showing L94 making contact with the GDIR motif, also showing proximity of the V1 loop (yellow) to the GDIR motif (orange) (**C**) and (**D**) CDRH3-V1 interactions and € Close-up of competition for the V3 residue D325 (thick orange loop) between PGT128 (orange) and 43A2 (cyan) (**F**) Overlay of 43A2 and non-human primate V1 loop binding polyclonal antibodies from animal 4O9, showing how both the rabbit monoclonal antibody and the NHP polyclonal antibodies avoid glycans in order to bind the N332 glycan supersite region on gp120(*31*).

**Fig. S4. Related to Figure 6.**
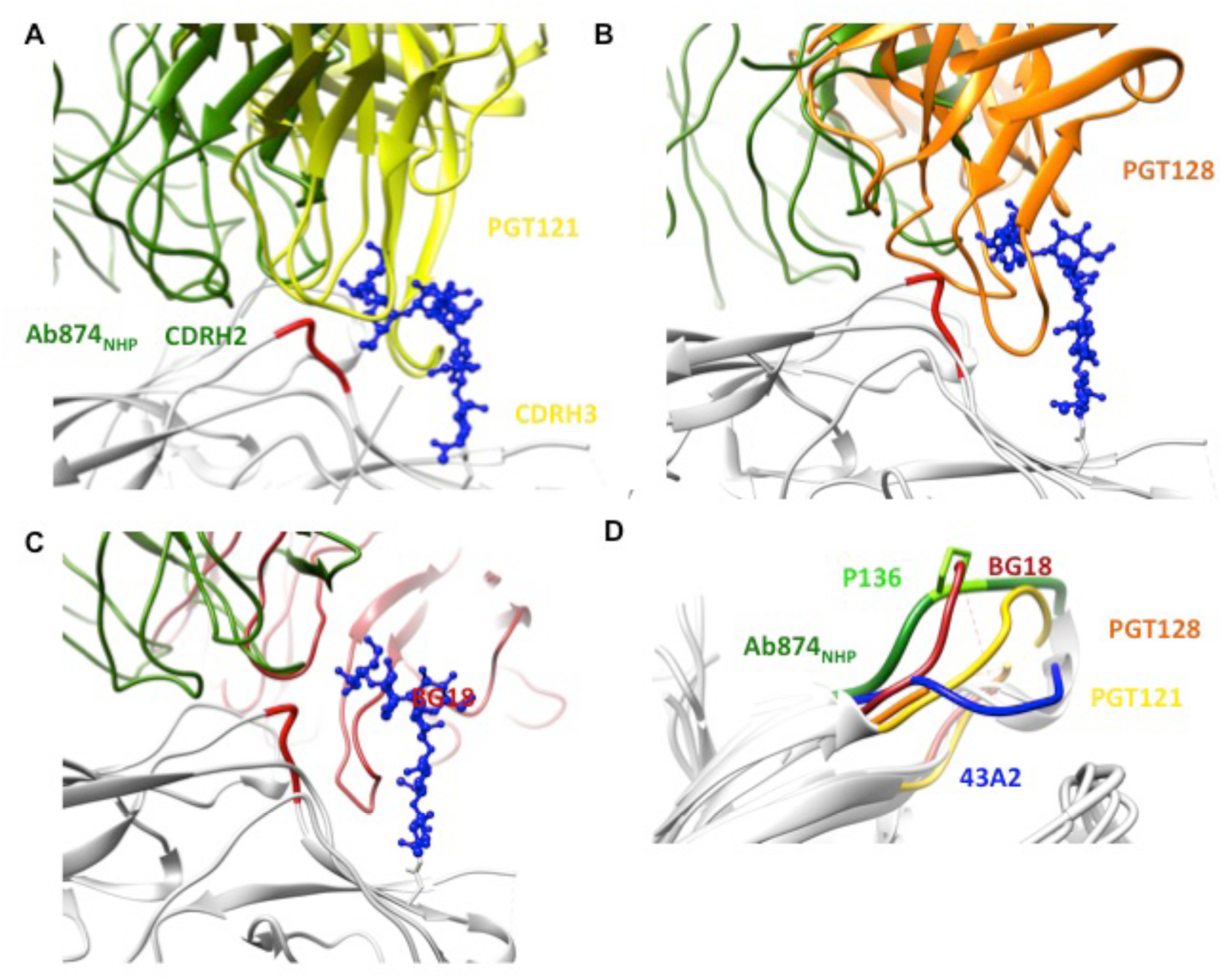
Comparison of N332 glycan interactions between bnAbs and Ab874_NHP_(*34*) (**A**), (**B**) and (**C**) Ab874_NHP_’s relatively short CDRH2 is positioned to potentially make minor contacts with an N332 glycan terminal mannose while bnAbs PGT121, PGT128, and B18, respectively, make both explicit and substantial contacts with both the N332 glycan and N332 (PGT121 and BG18) using their long CDH3’s, and (**D**) Comparison of V1 loop conformations in the ground state (43A2) and states induced by bnAbs or via a N136P mutation (Ab874_NHP_).

**Table S1.**
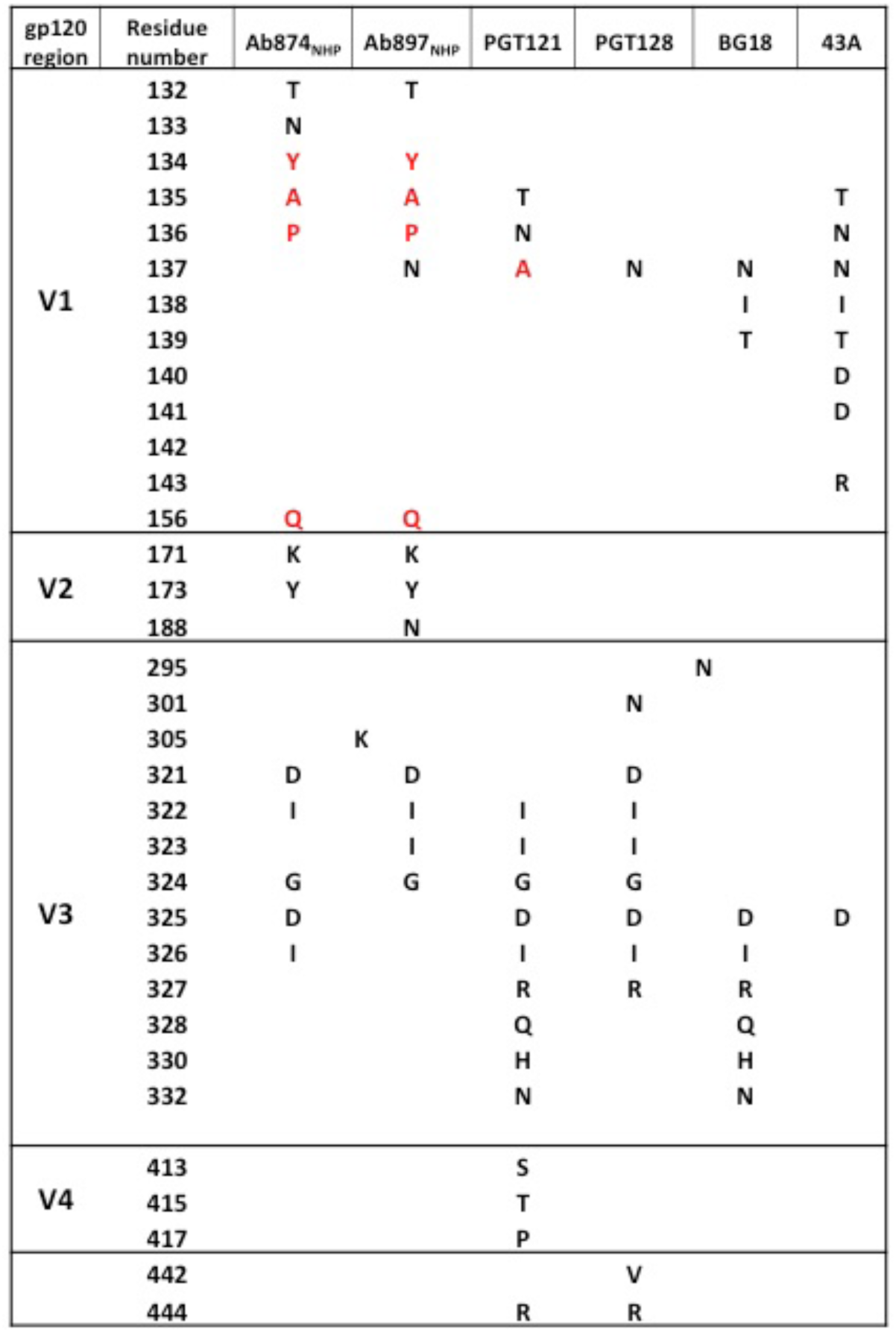
Ab874_NHP_, Ab897_NHP_, 43A2 BG505 epitope contacts vs N332 glycan supersite bnAbs. Antibody-gp120 side chain contacts (with radius < 4.0 A) of mAbs derived from immunized NHPs and rabbits primarily interact with the V1/V2 regions while predominant interactions by bnAbs are with the V3 base (based on PDB IDs: 6ORO, 6ORP, 5CEZ, 5ACO, 6CH7 corresponding to Ab874_NHP_, Ab897_NHP_, PGT121, PGT128, and BG18). Red type indicates non-native residues

**Table S2.** Related to Figures 5 and 6. CryoEM model building and statistics.

| Complex |  |  |
| --- | --- | --- |
| Composition (#) |  |  |
| Chains | 54 |  |
| Atoms | 19713 (Hydrogens: 0) |  |
| Residues | Protein: 2400 | Nucleotide: 0 |
| Water | 0 |  |
| Ligands | BMA: 3 NAG: 66 |  |
| Bonds (RMSD) |  |  |
| Length (Å) (# > 4σ) | 0.020 (42) |  |
| Angles (°) (# > 4σ) | 1.775 (338) |  |
| MolProbity score | 1.2 |  |
| Clash score | 1.65 |  |
| Ramachandran plot (%) |  |  |
| Outliers | 0.51 |  |
| Allowed | 3.57 |  |
| Favored | 95.92 |  |
| Rotamer outliers (%) | 0.14 |  |
| Cβ outliers (%) | 0 |  |
| Peptide plane (%) |  |  |
| Cis proline/general | 3.2/0.0 |  |
| Twisted proline/general | 0.0/0.0 |  |
| CaBLAM outliers (%) | 4.04 |  |
| ADP (B-factors) |  |  |
| Iso/Aniso (#) | 19713/0 |  |
| min/max/mean |  |  |
| Protein | 20.56/234.25/70.31 |  |
| Nucleotide | --- |  |
| Ligand | 41.66/121.29/78.68 |  |
| Water | --- |  |
| Occupancy |  |  |
| Mean | 1 |  |
| occ = 1 (%) | 100 |  |
| 0 < occ < 1 (%) | 0 |  |
| occ > 1 (%) | 0 |  |
| Data |  |  |
| Box |  |  |
| Lengths (Å) | 165.83, 163.77, 147.29 |  |
| Angles (°) | 90.00, 90.00, 90.00 |  |
| Supplied Resolution (Å) | 3.5 |  |
| Resolution Estimates (Å) | Masked | Unmasked |
| d FSC (half maps; 0.143) | --- | --- |
| d 99 (full/half1/half2) | 4.0/---/--- | 3.7/---/--- |
| d model | 3.9 | 3.9 |
| d FSC model<br>(0/0.143/0.5) | 3.4/3.5/3.7 | 3.5/3.5/4.1 |
| Map min/max/mean | -1.01/2.38/0.00 |  |
| Model vs. Data |  |  |
| CC (mask) | 0.83 |  |
| CC (box) | 0.77 |  |
| CC (peaks) | 0.72 |  |
| CC (volume) | 0.82 |  |
| Mean CC for ligands | 0.75 |  |
| EMRinger: 2.58 |  |  |
| CARP: pass |  |  |
| pdbe-care: pass |  |  |
| Privateer: pass |  |  |

